# *rde-3* reduces piRNA-mediated silencing and abolishes inherited silencing in *C. elegans*

**DOI:** 10.1101/2022.08.13.503839

**Authors:** Monika Priyadarshini, Sarah AlHarbi, Christian Frøkjær-Jensen

## Abstract

Small RNA-mediated silencing of target genes can persist across generations and *C. elegans* is a well-established model for studying the molecular basis for epigenetic inheritance. We recently developed a piRNA-based inherited silencing assay that causes a high incidence of males by targeting *him-5* and *him-8*. Acute gene silencing is determined in the presence of the piRNAi extra-chromosomal array and inherited silencing after loss of the piRNA trigger. This assay has the advantage of targeting endogenous genes that are easily scored in mutant backgrounds and obviates the need for mutant validation and genetic crosses, which can influence inherited silencing. Here we show an example of the assay by testing acute and inherited piRNA-mediated *him-5* silencing in ribonucleotidyltransferase *rde-3* (*ne3370*) mutant animals. In the absence of *rde-3*, acute silencing was reduced but still detectable, whereas inherited silencing was abolished.

---

*C. elegans* is a convenient model for studying small RNA-mediated inherited silencing due to the animal’s short generation time (three days) and the ability to identify molecular pathways in genetic screens^1–3^. Epigenetic silencing of an endogenous gene is often done by targeting a temperature-sensitive gain-of-function allele of *oma-1(zu405)* with dsRNA and silencing persists for up to three generations^4^. For a visual read-out, single-copy transgenes with GFP expression in the germline^5–7^ have been engineered to contain endogenous piRNA binding sites in the 3’ UTR^8–11^. For transgenes, piRNA-induced silencing persists longer and sometimes indefinitely. Genetic factors required for small RNA-mediated inherited silencing have primarily been identified by crossing silenced piRNA *gfp* sensor strains into genetic mutant backgrounds^8,10–12^. However, introducing mutations by genetic crosses raises several concerns. First, there are several examples of mating causing changes in epigenetic inheritance. For example, the lack of transgene pairing during meiosis after a cross can lead to permanent transgene silencing via PRG-1-dependent mechanisms^13^and mating can induce multigenerational silencing inherited for over 300 generations^14^. Moreover, Dodson and Kennedy^15^ characterized a transgenerational disconnect between the genotype and phenotype (sensitivity to exogenous RNAi) of *meg-3/4* mutants for more than seven generations after a genetic cross. Second, crosses frequently require molecular genotyping which makes it cumbersome to perform many biological replicates. Third, there are some concerns about using transgenes as a proxy for endogenous gene silencing. For example, most piRNA sensor strains include synthetic piRNA binding sites in the 3’ UTR^8–11^ but endogenous genes are resistant to piRNA silencing when targeting their 3’ UTRs^16,17^. Moreover, transgene insertion site^18^, noncoding DNA structures^19^, coding sequence^20,21^, and transgene structure^22^ can influence epigenetic silencing. These observations suggest that transgenes may not fully recapitulate the balance between silencing foreign DNA and protecting endogenous gene expression^23^. Finally, distinguishing between silencing initiation and maintenance phases is complicated using genetic crosses. Experiments require crossing mutant allele to sensor strains, de-repress silencing, and outcrossing mutations to monitor *de novo* establishment of silencing^11^.

We recently developed a method called piRNA interference (piRNAi) that can efficiently silence both transgenes and endogenous genes by expressing synthetic piRNAs from arrays generated by injection^16,24^. Using piRNAi, we identified two endogenous targets, *him-5* and *him-8*, that inherit silencing for three and six generations, respectively^16^. *him-5^25^* and *him-8^26^* mutants are generally healthy but have a similar loss-of-function phenotype that is easy to score (~35% males in the population). We reasoned that piRNA-mediated silencing of *him-5* or *him-8* might be useful as a tool to directly test the role of gene mutations in initiating and maintaining inherited silencing. Here, we show that piRNAi can be used to test acute and inherited silencing in *rde-3* (also known as *mut-2*^27^).

## Results and discussion

*rde-3* is required for Tc1 transposon silencing in the germline^28^ and RNA interference (RNAi)^29^. *In vitro*, RDE-3 has ribonucleotidyltransferase activity^30^ and, in *vivo*, *rde-3* is required for the addition of non-templated poly (UG) tails to the 3’ end of mRNAs targeted by RNAi and repressed transposons^31^. pUGylated mRNAs are templates for RNA-dependent RNA polymerases (RdRPs) resulting in small RNA amplification and inherited silencing^31^. RDE-3 is required to maintain silencing of piRNA transgene sensors^9–11^. However, the role of *rde-3* in initiating silencing is not clear; re-introducing RDE-3 led to rapid re-silencing of a *gfp::cdk-1* transgene but variable and incomplete re-silencing of a *gfp*::*csr-1* transgene^11^. These conflicting results could be caused by differences between transgenes or by effects of mating. We, therefore, decided to use piRNAi to test the role of RDE-3 in initiation and maintenance of silencing of an endogenous gene. We targeted *him-5* with six synthetic guide piRNAs (sg-piRNAs) (**Fig. 1A-B**) in wildtype (N2) animals and *rde-3(ne3370)* mutants. *rde-3* is a mutator strain and is relatively unhealthy, with a small brood size and infrequently produces males. To account for an elevated male frequency in the mutant population, we generated transgenic *rde-3* animals with non-targeting sg-piRNAs as a control. Targeting *him-5* with piRNAi resulted in an increased frequency of males in N2 animals but a significantly lower male frequency in *rde-3* animals (30 ± 1.9% vs 7.6 ± 1.3%, P = 0.05, mean ± SEM) (**Fig. 1C**). However, male frequency in *rde-3* animals was significantly increased compared to negative controls (7.6 ± 1.3% vs 0.2 ± 0.2%, P = 0.05, mean ± SEM). We tested the role of RDE-3 in maintaining silencing by losing the piRNAi trigger (the piRNAi arrays with a *Pmyo-2::mCherry* fluorescent marker) and scoring male frequency in the following generations. In wildtype animals, male frequency remains elevated for at least three generations after the primary piRNAs targeting *him-5* are lost (**Fig. 1D**), consistent with prior observations^16^. In contrast, we could not detect an inherited elevation of male frequency in *rde-3* mutants (**Fig. 1D**). The initial frequency of males was relatively low in *rde-3* animals which limits our ability to make strong conclusions. However, our results support a model where primary piRNAs can post-transcriptionally silence a target transcript *(him-5* mRNA) at reduced efficiency but *rde-3* is required for small RNA amplification and transcriptional silencing. Presumably, RDE-3 amplifies the primary trigger by generating *him-5* pUG RNAs that are templates for RdRP-mediated 22G amplification; these secondary RNAs are subsequently used to set up transcriptional silencing via repressive chromatin marks deposited by the *hrde-1* dependent nuclear RNAi pathway.

**Figure 1.**
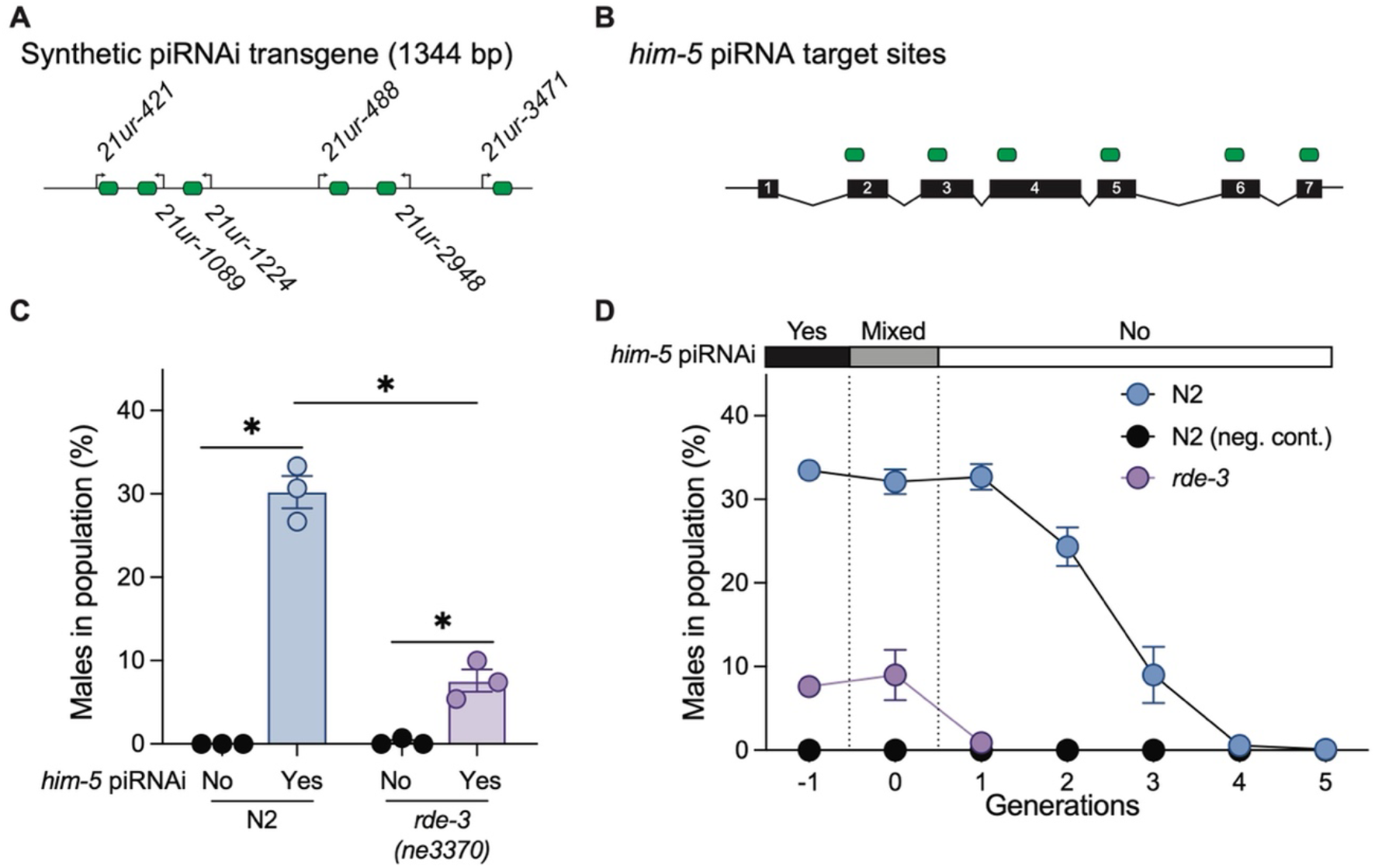
Testing acute and inherited silencing in *rde-3* mutants. **A**. Schematic of synthetic piRNAi construct. **B**. Synthetic piRNAs target sites in *him-5*. **C**. Quantification of males frequency after piRNAi against *him-5* in N2 wild type and *rde-3(ne3370)* animals; N2: N=3, *rde-3:* N=3 *(him-5* piRNAi and neg. control); Statistics: Mann-Whitney one-tailed test, N2 vs control (P = 0.05), *rde-3* vs control (P = 0.05), N2 vs *rde-3* (P = 0.05). **D**. Inherited *him-5* piRNAi silencing in N2 and *rde-3* animals; N = 3 all conditions. Negative control: non-targeting piRNAs. Bar above graph indicates the presence (“yes”), mixed generation (“mixed”), and absence (“no”) of a *Pmyo-2::mCherry* marked extra-chromosomal array expressing synthetic piRNAs targeting *him-5*. Each data point is a biologically independent transgenic line for panel C and the average of three biologically independent transgenic lines for panels D. Error bars indicate standard error of mean (SEM).

More generally, we demonstrate that piRNAi can be used as a tool to directly test genetic factors required for acute and inherited silencing of endogenous genes. Elevated male frequency (induced by targeting *him-5* or *him-8)* is easy to score in a variety of genetic backgrounds and allows distinguishing between silencing initiation and maintenance of endogenous genes.

## Acknowledgements

We thank the CGC, which is funded by the NIH Office of Research Infrastructure Programs (grant no. P40 OD010440) and WormBase for data and literature curation.

## Author Contributions

Monika Priyadarshini: Investigation, conceptualization, and the original draft.

Sarah Alharbi: Investigation, writing-review.

Christian Frøkjær-Jensen: Funding acquisition, conceptualization, writing-review, and editing.

Funding acquisition: CFJ

## Competing interests

The authors declare no competing interests.

## Materials & Methods

### Transgenesis

Transgenic animals with piRNAi extrachromosomal arrays were generated according to standard injection protocols ^32^. The injection mix for all experiments consisted of ~15-20 ng/μl of synthetic dsDNA piRNA fragments (Twist Bioscience), 12.5 ng/μl of a plasmid encoding hygromycin resistance (pCFJ782), and 2 ng/μl of a fluorescent co-injection marker P*myo-2::*mCherry (pCFJ90). The total concentration of the injection mix was adjusted to 100 ng/μl with 1kb DNA ladder (1 kb Plus DNA Ladder, catalog no. 10787018, Life Technologies). This mix was injected into young adult hermaphrodite animals and allowed to recover on standard NGM plates seed with OP50 bacteria. 36-48 hours post-injection, 500 μl of 4 mg/ml stock of Hygromycin solution (Gold Biotechnology, catalog no. H-270-1) was topically added to the bacterial lawn of injection plates to select for transgenic (F1) progeny. A single healthy transgenic F2 adult was picked from each plate to generate a clonal strain and pharyngeal mCherry fluorescence was visually confirmed.

### Quantification of male frequency

Quantification of male frequency was performed as previously reported by ^16^. Briefly, six virgin L4 hermaphrodites were picked to freshly seeded NGM plates with hygromycin selection to select for the piRNAi array. The frequency of males was determined using a dissection microscope and by visual inspection of 100 adult animals on plates incubated on ice for 30 minutes to immobilize animals.

### Inherited silencing assay

Six virgin L4 animals were picked to non-selective NGM plates to obtain a mixed progeny population with and without the piRNAi array. Males were not quantified in this mixed population; however, L4 animals carrying the sg-piRNAs were propagated in parallel, and their progeny were scored for males (G0). In the following generation, non-transgenic L4 animals were carefully picked from the mixed population based on the absence of pharyngeal mCherry expression (a marker for the piRNAi array). The progeny of these animals was quantified for male frequency (G1). Male frequency was quantified in all following generations by picking L4s until the male frequency was below 1%.

### Data quantification and statistics

Independently generated transgenic animals were treated as biological replicates. piRNA-mediated silencing is stochastic (i.e., most strains show robust silencing but some strains are not silenced at all) and the data do not follow a normal distribution. We performed statistical tests using a one-side parametric Mann-Whitney tests to account for this.

### Software

Statistical analysis was performed with GraphPad Prism (v 9.4.1), figures were generated with Adobe Illustrator (v 26.4.1) and the manuscript was written with Microsoft Word (v 16.63.1).

#### Reagents

<u>List of piRNAi fragments, strains, and plasmids used in this study</u>

#### Strains

N2 Standard wiltype strain^33^

WM286 *rde-3(ne3370)* I^34^

#### Plasmids and transgenes

pCFJ90 *Pmyo-2::mCherry::unc-54* UTR^35^

pCFJ782 P*rps-0::*HygroR

##### *him-5* (six targeting piRNAs in upper-case)

cgcgcttgacgcgctagtcaactaacataaaaaaggtgaaacattgcgaggatacatagaaaaaacaatacttcgaattcatttttcaattacaaatcctgaaatgtttcactgtgttcctataagaaaacattgaaacaaaatattaagTGAGTTAGCTTTCCGGAGCTTctaattttgattttgattttgaaatcgaatttgcaaatccaattaaaaatcattttctgataattagacagttccttatcgttaattttattatatctatcgagttagaaattgcaacgaagataatgtcttccaaatactgaaaatttgaaaatatgttTCCTCACGAAAAACCTGCCTAttGccagaactcaaaatatgaaatttttatagttttgttgaaacagtaagaaaatcttgtaattactgtaaactgtttgctttttttaaagtcaacctacttcaaatctacttcaaaaattataatgtttcaaattacataactgtgtATGCAGAGAGATCAGTAGGTActgtagagcttcaatgttgataagatttattaacacagtgaaacaggtaatagttgtttgttgcaaaatcggaaatctctacatttcatatggtttttaattacaggtttgttttataaaataattgtgtgatggatattattttcagacctcatactaatctgcaaaccttcaaacaatatgtgaagtctactctgtttcactcaaccattcatttcaatttggaaaaaaatcaaagaaatgttgaaaaattttcctgtttcaacattatgacaaaaatgttatgattttaataaaaacaaTCGATCACTGTTGACAATCACttctgtttttcttagaagtgttttccggaaacgcgtaattggttttatcacaaatcgaaaacaaacaaaaatttttttaattatttctttgctagttttgtagttgaaaattcactataatcatgaataagtgagctgcccaagtaaacaaagaaaatttggcagcggccgacaactaccgggttgcccgatttatcagtggaggaTAATCCGGCACGTAGAATGTAtctaatgtgatgtacacggttttcatttaaaaacaaattgaaacagaaatgactacattttcaaattgtctatttttgctgtgtttattttgccaccaacaaTCGATGCGACCAACTGTTTTTtcaatctagtaaactcacttaatgcaattcctccagccacatatgtaaacgttgtatacatgcagaaaacggttttttggttttaatgggaacttttgacaaattgttcgaaaatcttaagctgtcccatttcagttgggtgatcgattt

##### Control (non-targeting piRNAs)

ctcggtcaattaaagaaagaCATTTTTCATCGGATTTGCTActaaaaaataattttaaAAACGATCATATGCAAATCCAgtgaaactttattcaaaccaaaacgtttaatcagctaattgaaacattaaaaattttatgattttgttagtttttctagcaatgtcaatgcaatcaaataattttcaagtaagatgtttaatgagttatagacttttttattaaatttttgaaaaaaaaaccgatttcagatttaagtaaaattatctctgcttctgctgcattgctgcgaaacaaaaattcctttctgtgcaaagtatagtATAAACGAGGAGCACAAATGAgtgacaattagaaatctcaccgggttttctagatcatctgaaacatataattttaaaaaattgacaccttgttcaacTGTTGCACATATCACTTTTGAtcgaaacattaaatgtctcatgatttttaaagctcttttagaacagtcgCCAATCCCCTTATCCAATTTAttgaaaacaattttctagcgagatgttaaatgagtttgttgaaacagtagattttcgtgtaaacttttgaaaacaaaaattacgttttaaataaaattatatccacttcagcagtgtgcccttgaaacaaaaaagctcgatcaaaaaatttattttttgtgaatggccaccaacttttcaggcaaaattacaaaaaaacataaaatttactgtttcaaaaagttaatataattttggcagcgcatatacctacacTGAATTTTGGCAGAGGCAATTacctctttttgaaaataaag

## Notes

### Competing Interest Statement

The authors have declared no competing interest.

